# Identification of a new ORF3a-E fusion subgenomic RNA of SARS-CoV-2 and its biological features in the infection process

**DOI:** 10.1101/2024.08.28.610093

**Authors:** Yifan Zhang, Huiwen Zheng, Jing Li, Xinglong Zhang, Xin Zhang, Jiali Li, Heng Li, Xin Zhao, Zihan Zhang, Yingyan Li, Keqi Chen, Shasha Peng, Ying Zhang, Haijing Shi, Longding Liu

## Abstract

Subgenomic RNAs (sgRNAs) are discontinuous transcription products of severe acute respiratory syndrome coronavirus 2 (SARS-CoV-2) that are involved in viral gene expression and replication, but their exact functions are still being studied. Here, we report the identification of a new type of sgRNA, the fusion ORF3a-E-sgRNA, involved in the infection process of SARS-CoV-2. This sgRNA codes both ORF3a and E and can be detected throughout the viral life cycle in SARS-CoV-2-infected cells with high copy numbers. ORF3a-E-sgRNA guides more ORF3a translation and promotes the expression of cellular ribosomal protein S3 (RPS3) and the binding of eukaryotic translation initiation factor 4E (eIF4E). Single-cell sequencing of a SARS-CoV-2-infected human bronchial epithelial cell line (16HBE) revealed that maintenance of this stable translational environment by ORF3a-E-sgRNA is important for the SARS-CoV-2 assembly and release capabilities and is also beneficial for viral evasion of host innate immunity. More importantly, the transcription level of ORF3a-E-sgRNA contributes to differences in infection processes between the Wuhan strain and XBB strain of SARS-CoV-2.

## Introduction

Subgenomic RNA (sgRNA) is produced by discontinuous transcription from genomic RNA via the transcription complex (RTC) and the transcription replication complex (TRS). The SARS-CoV-2 genome contains 14 open reading frames (ORFs) that are preceded by the TRS body sequence (TRS-B). Additionally, there is a TRS leader sequence (TRS-L) located after the conserved 5′ leader sequence, with a core ACGAAC sequence that can be used for sgRNA detection. When the RTC encounters TRS-B, it performs discontinuous transcription with TRS-L, generating different sgRNAs of different sizes^1-3^. To date, nine types of sgRNAs have been clearly identified, including S, ORF3a, E, M, ORF6, ORF7, ORF8, N and ORF10^4,5^. However, owing to the joining of the 5’ and 3’ positions of the leader sequence, long-distance fusion independent of TRS-L, and local fusion causing deletions or additions in structural and accessory genes, novel sgRNA types may also exist during SARS-CoV-2 infection.

sgRNAs and their encoded proteins are involved throughout the viral lifecycle, from infection to release. This is closely related to the ability of SARS-CoV-2 to rapidly replicate during infection, providing guidance for virus synthesis and helping the virus assemble into complete viral particles. The close proximity of TRS-L and TRS-B sequences during discontinuous transcription increases the recombination frequency, increasing the synthesis speed of sgRNAs and aiding virus functions^4,6^. There are four types of sgRNAs, encoding the spike protein (S), nucleocapsid protein (N), membrane protein (M), and envelope protein (E), which encapsulate genomic RNA (gRNA) to form new viral particles that are released by cells. The remaining sgRNAs encode viral accessory proteins, which are a highly variable set of virus-specific proteins that help regulate the host response to infection. The sgRNAs of SARS-CoV-2 can assist in viral evasion of host immune defenses during infection, such as N, ORF9b, and ORF6, which influence the transcription of IFN-stimulated genes (ISGs) and accelerate viral replication via an increase in the corresponding RNA and protein levels^7-9^. Upon infection, SARS-CoV-2 may impact the host internal translation environment, usually by inhibiting the host’s translation activity, and the sgRNAs continue to initiate translation because of the presence of a leader sequence in the 5’ UTR.

SARS-CoV-2 variants have led to multiple waves of infection worldwide, such as the Alpha (B.1.1.7) variant from the UK in September 2020, the Beta (B.1.351) variant from South Africa in October 2020, the Gamma (P.1) variant from Brazil in November 2020, the Delta (B.1.617.2) variant from India in April 2021, and the Omicron (B.1.1.529) variant, which has been circulating since November 2021.

Research indicates that as these variants evolve, they undergo significant changes in replication capacity, pathogenicity, and transmissibility^10^. For example, in transgenic mouse and hamster models with human ACE2 receptors, the Alpha and Delta variants exhibited extensive replication across multiple organs. Moreover, the Omicron variant, although more rapidly spreading than the Delta variant, shows substantially reduced infectivity and pathogenicity^11-13^. Several factors contribute to the varying transmission and replication capabilities among these variants. The variants can enhance their ability to enter cells by improving fusion with the host cell membrane, better interacting with ACE2 receptors of other species, enabling cross-species transmission, and partially evading preexisting immunity within populations^14^. Furthermore, variations in sgRNA frequency, the proteins they encode, and host cell activity levels can limit the resources available for viral particle formation, thus reducing replication capabilities. Studies suggest that infections with the Alpha variant have increased sgRNA abundance, which is closely tied to host cellular activities^15^. Therefore, the roles and functions of sgRNAs are critical during SARS-CoV-2 replication and significantly influence viral replication efficiency.

In our study, we investigated the discontinuous transcription of SARS-CoV-2 sgRNAs and discovered a novel sgRNA, in addition to the nine previously identified, which we named ORF3a-E-sgRNA. This new RNA includes a fusion of the sequences of both ORF3a and E, enabling it to encode both the structural protein E and the nonstructural protein ORF3a simultaneously. Our results indicate that ORF3a-E-sgRNA notably recruits more of the cellular ribosomal protein S3 (RPS3) and binds more effectively with the eukaryotic translation initiation factor 4E (eIF4E) at the 5’ UTR cap, thereby increasing local translation levels. This leads to increased expression of the ORF3a protein, which subsequently impacts the replication ability of SARS-CoV-2. Notably, we observed significant differences in virus replication and host translation levels between 16HBE cells infected with the original 2019 Wuhan strain and those infected with the 2022 Omicron variant. Additionally, ORF3a-E-sgRNA seems to impair the host antiviral immune response to some extent, fostering a more favorable environment for SARS-CoV-2 replication.

## Results

### 1. Identification of ORF3a-E-sgRNA during SARS-CoV-2 infection

Using primers that bind between the 5’ UTR, the leader core sequence, and the sg-ORF, we investigated various sgRNAs, such as M-sgRNA, S-sgRNA, N-sgRNA, ORF7a-sgRNA, ORF8-sgRNA, and E-sgRNA (Figure S1A-B). Our analysis revealed a common sequence structure: 5’ UTR (75 bp (ACGAAC) + variable RNA fragment) - ORF - 3’ UTR. While most of the PCR fragment sizes matched the expected values, we observed an unexpected strong signal for E-sgRNA that was approximately 800 bp larger than predicted. Sequencing confirmed this as a fusion sgRNA linking ORF3a with E, resulting in the structure 5’ UTR (75 bp (ACGAAC) + variable RNA fragment) - ORF3a - ORF - medi-E - ORF - 3’ UTR (Figure 1A). We named this novel sgRNA ORF3a-E-sgRNA. From the result of 5’-RACE of ORF3a-E-sgRNA, we validated its complete structure without extraneous sequences at the 5’ end (Figure S1C). For detecting this molecule accurately, a 3’ fluorescence-labeled probe was designed to target segments of ORF3a-E-sgRNA, and its specificity and sensitivity were confirmed (Figure S1D-F). The synthesized ORF3a-E-sgmRNA was capped, polyadenylated, and transfected into 293T cells. Fluorescence probing and FLIM-FRET were employed to measure sequence distances, revealing a decrease in the fluorescence lifetime by approximately 0.2 ns upon the addition of the 3a linker E probe (Figure 1D-E). This result was also observed when ORF3a-E-sgRNA was detected with another pair of probes (Figure 1F). By using this probe, we confirmed ORF3a-E-sgRNA in different cell lines, such as 16HBE and AC16 cells infected with SARS-CoV-2 from Wuhan and XBB at an M.O.I. of 0.1, as well as in the heart and kidney tissues of SARS-CoV-2-infected rhesus monkeys (Figure 1G-H). Single-cell sequencing revealed a positive correlation between ORF3a-E-sgRNA and other sgRNAs, particularly ORF3a-sgRNA and E-sgRNA (Figure 1I-K). Thus, our research confirmed that ORF3a-E-sgRNA, a novel sgRNA with unique sequence characteristics, is produced during SARS-CoV-2 infection.

**Figure 1.**
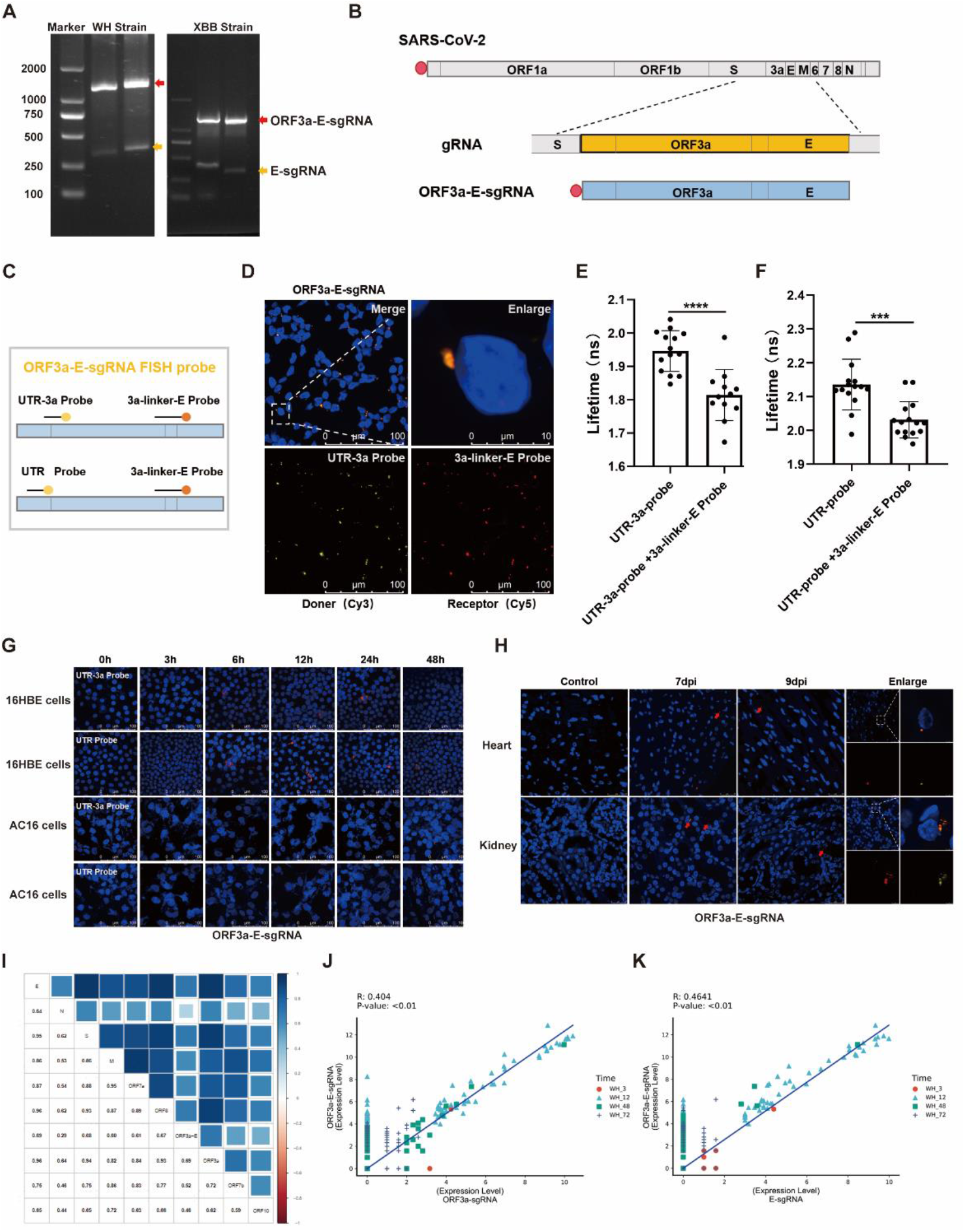
Detection and validation of ORF3a-E-sgRNA during SARS-CoV-2 infection. (A) Detection of ORF3a-E-sgRNA in 16HBE and AC16 cells infected with the Wuhan and XBB strains. (B) Schematic diagram of the sequence characteristics of ORF3a-E-sgRNA. (C) Schematic diagram of probe design targeting the ORF3a-E-sgRNA sequence. (D) Analysis of ORF3a-E-sgRNA-transfected 293T cells using a labeled probe for in situ hybridization; a partially enlarged image is shown. (E-F) Fluorescence resonance energy transfer was used to calculate the fluorescence lifetime of UTR-3a probes (E) or UTR probes (F) with or without 3a-linker-E probes for labeling ORF3a-E-sgRNA. (G) Detection of ORF3a-E-sgRNA within 48 hours of SARS-CoV-2 infection in 16HBE and AC16 cells at an M.O.I. of 0.1. (H) Detection of ORF3a-E-sgRNA in the heart and kidney tissues of rhesus monkeys on the seventh and ninth days after SARS-CoV-2infection. (I) Analysis of the correlation between ORF3a-E-sgRNA and other sgRNAs. (J-K) Analysis of the correlation between ORF3a-E-sgRNA and ORF3a-sgRNA (J) or E-sgRNA(K).

### 2 ORF3a-E-sgRNA increases ribosomal subunit protein accessibility and improves the translation efficiency of ORF3a

The SARS-CoV-2 sgRNA is capped, indicating that its protein translation occurs through a cap-dependent mechanism involving multiple eukaryotic initiation factors, such as the eIF4F complex. To explore the functions of ORF3a-E-sgRNA, we generated this sgRNA via in vitro transcription, capping, and tailing. Immunoprecipitation (IP) and RNA pull-down experiments revealed that ORF3a-E-sgRNA, similar to the ORF3a-only sgRNA, binds to most subgenomic interacting proteins. However, the ORF3a-E-sgRNA specifically binds more strongly to the cap-binding eukaryotic translation initiation factor 4E (eIF4E). This increased eIF4E binding leads to increased expression of the ORF3a protein and greater recruitment of ribosomes, as indicated by the increase in the levels of the ribosomal protein RPS3 (Figure 2A-D). Fluorescence imaging confirmed that ORF3a-E-sgRNA induces early and sustained expression of eIF4E (Figure S2). Compared to transfection with ORF3a-sgRNA alone, transfection with ORF3a-E-sgRNA alone or cotransfection with ORF3a-sgRNA and ORF3a-E-sgRNA significantly increased RPS3 levels in host cells (Figure 2E-F). These findings suggest that ORF3a-E-sgRNA has a relatively strong ability to recruit ribosomes and translation initiation factors. To further investigate the impact of this enhanced ribosome recruitment, we coexpressed an exogenous reporter gene with the different sgRNAs. Both western blot and fluorescence analyses revealed that the translation of the reporter gene was more efficient when it was coexpressed with ORF3a-E-sgRNA than when it was coexpressed with ORF3a-sgRNA (Figure 2G-H). Moreover, we found that ORF3a-E, similar to ORF3a (as reported in the literature^16^), can promote lysosomal exocytosis, leading to increased release of virus particles from infected cells. This was demonstrated by the increased levels of the lysosomal membrane protein LAMP1 on the cell surface (Figure 2I). The above results indicate that compared with the ORF3a-only sgRNA, the ORF3a-E-sgRNA has a greater ability to impact the host cell translation machinery, supporting increased expression of the ORF3a protein and viral particle release.

**Figure 2.**
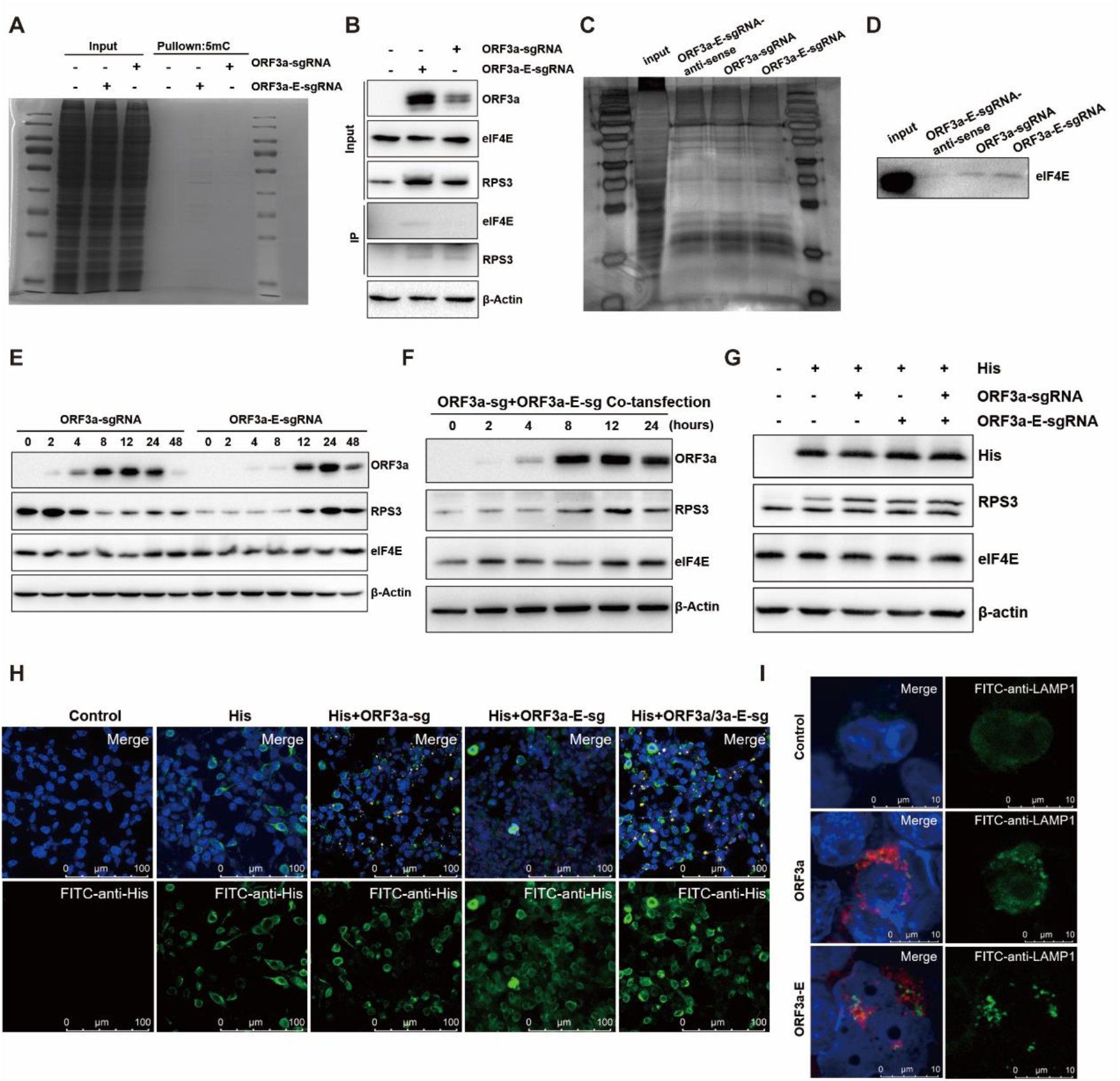
Functional analysis of proteins encoded by ORF3a-E-sgRNA. (A) Coomassie Brilliant Blue staining of the gel after immunoprecipitation of cells transfected with ORF3a-sgRNA and ORF3a-E-sgRNA. (B) The binding of ORF3a-sgRNA and ORF3a-E-sgRNA to ribosomal proteins and translation initiation factors in 293T cells after immunoprecipitation. (C) Silver staining was performed to detect ORF3a-sgRNA- and ORF3a-E-sgRNA-bound proteins from RNA pull-down experiments. (D) Detection of the binding of ORF3a-sgRNA and ORF3a-E-sgRNA with translation initiation factor eIF4E. (E) Detection of ribosome-associated proteins in 293T cells transfected with ORF3a-sgRNA or ORF3a-E-sgRNA. (F) Detection of ribosome-associated proteins in 293T cells cotransfected with ORF3a-sgRNA and ORF3a-E-sgRNA. (G) Detection of the expression of his in 293T cells with or without sgRNA transfection. (H) Detection of the fluorescence intensity of his in 293T cells cotransfected with his and sgRNA. (I) Expression of LAMP1 on the cell membrane surface after the expression of ORF3a-sgRNA and ORF3a-E-sgRNA.

### 3 SARS-CoV-2 variant showed different replication processes related to ORF3a-E-sgRNA

We examined the functional characteristics of ORF3a-E-sgRNA during SARS-CoV-2 infection. Samples from 16HBE cells were collected at various intervals post infection with the 2019 Wuhan prototype strain and the 2022 Omicron variant strain XBB, both at an M.O.I. of 0.1. Single-cell sequencing captured different sgRNA types, including ORF3a-E-sgRNA. Q-PCR and sc-RNA-seq results revealed that viral loads increased consistently in both 16HBE cells following infection with the Wuhan strain of SARS-CoV-2, whereas the XBB strain’s viral load remained stable over 72 hours. The viral load generated by the Wuhan strain significantly exceeded that of the XBB strain (Figure 3A-B). Moreover, differential gene expression and pathway enrichment analyses revealed that the Wuhan strain maintained a replicative state from the initial to final infection stages, mobilizing ribosomes and the endoplasmic reticulum within the host. In contrast, XBB strain infection significantly downregulated ribosome-related pathways, potentially due to ORF3a-E-sgRNA (Figure S3, S4).

**Figure 3:**
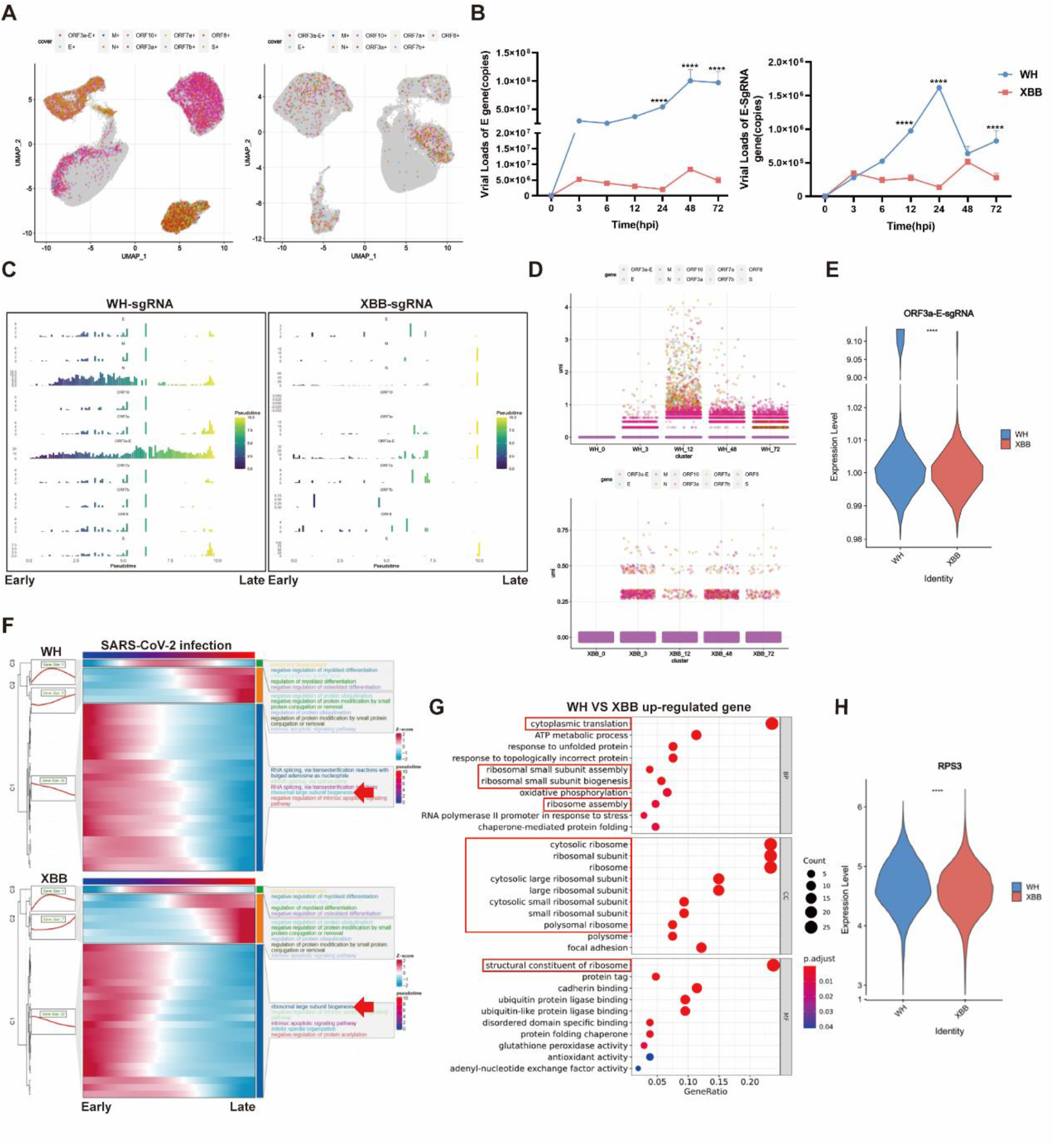
Single-cell sequencing analysis of the functional characteristics of ORF3a-E-sgRNA in different variants of SARS-CoV-2. (A) Annotation of viral particles in 16HBE cells infected with the Wuhan and XBB strains. (B) Detection of E-gRNA and E-sgRNA in 16HBE cells infected with the Wuhan and XBB strains via Q-PCR. (C) Changes in the proportion of sgRNA in 16HBE cells infected with the Wuhan or XBB strain over pseudotime. (D) Changes in the proportion of sgRNA in 16HBE cells infected with the Wuhan or XBB strain over the infection period. (E) Comparison of the expression ratios of ORF3a-E-sgRNA in Wuhan strain- and XBB strain-infected 16HBE cells. (F) Differential pathway enrichment in 16HBE cells infected with the Wuhan or XBB strain over pseudotime. (G) Comparison of host differentially expressed gene pathway enrichment in 16HBE cells infected with the Wuhan or XBB strain. (H) Comparison of the expression levels of RPS3 in 16HBE cells infected with the Wuhan or XBB strain.

We further analyzed the dynamics of SARS-CoV-2 sgRNAs during infection via both pseudotime and infection timing approaches. The pseudotime analysis simulated the SARS-CoV-2 infection cycle on the basis of changes in gene expression during different stages of infection, dividing it into early, middle, and late stages. We annotated the dynamic changes in different sgRNA types as the pseudoinfection time varied (Figure S5). The results revealed that the expression proportion of Wuhan sgRNAs dynamically changed as pseudotime progressed, with their distribution being relatively concentrated in the first half of the pseudoperiod. Among them, ORF3a-E-sgRNA was present throughout the entire pseudoinfection period. In contrast, the distribution of XBB sgRNA is scattered and lacked an obvious pattern (Figure 3C). The infection timing results further revealed that ORF3a-E-sgRNA was most abundant among all the sgRNAs during Wuhan strain infection, indicating its active role in progeny virus formation. However, this ORF3a-E-sgRNA activity was not detected after XBB strain infection (Figure 3D-E).

We further investigated the functional effects of ORF3a-E-sgRNA in different SARS-CoV-2 variants. The host gene expression patterns were similar between Wuhan and XBB strain infections in terms of both the infection time and pseudotime series. Notably, ribosomal functional activity was high in the first half of the pseudoperiod, which coincided with the expression pattern of ORF3a-E-sgRNA (Figure 3F). Targeted analysis of sgRNAs using sc-RNA and functional enrichment revealed that most sgRNAs were not enriched in host pathways during XBB strain infection compared to Wuhan strain infection. However, the ORF3a-E-sgRNA from the Wuhan strain was enriched in pathways related to protein synthesis, such as protein folding, ER-Golgi transport, and protein stability (Figure S6-S8). Furthermore, the ribosome-related gene set was highly enriched in hosts infected with the Wuhan strain, and the translation process marked by the 40S ribosomal protein RPS3 was significantly upregulated (Figure 3G-H). Overall, the activity of ORF3a-E-sgRNA appears to be a key factor underlying the differences in host protein synthesis between SARS-CoV-2 variants.

### 4 ORF3a-E-sgRNA limits the innate immune response for effective replication of SARS-CoV-2

The structural and nonstructural proteins of SARS-CoV-2 play crucial roles in viral evasion the host antiviral immune response, primarily by targeting key molecules in the IFN-I signaling pathway. This process may also involve ORF3a-E-sgRNA and its high expression during SARS-CoV-2 infection. We observed that in 16HBE cells infected with the Wuhan or XBB strain, the expression of interferon-beta (IFN-β) and the interferon-stimulated gene ISG15 remained low within the first 48 hours but then rapidly increased at 72 hours, potentially leading to an inflammatory cytokine storm (Figure 4A-B). Further analysis revealed that the expression of many interferon-stimulated genes increased significantly as the infection progressed, especially at 72 hours (Figure 4C). This inverse correlation between sgRNA expression and the immune response suggests the presence of innate immune antagonists, such as ORF6 and ORF9b. To verify the immunity evasion ability of ORF3a-E-sgRNA, we transfected it into 293T cells. The results revealed that IFN-α, IFN-β, and IFN-γ expression increased at certain transfection doses but decreased at higher doses (Figure 4D). Importantly, compared with ORF3a-sgRNA alone, ORF3a-E-sgRNA exhibited stronger immune evasion ability (Figure 4E-F). Moreover, the ORF3a-E-sgRNA level was positively correlated with genomic RNA levels but negatively correlated with the expression of the interferon-stimulated gene ISG15 (Figure 4G-H). In summary, ORF3a-E-sgRNA helps SARS-CoV-2 maintain low host innate immunity during its life cycle, enabling effective viral replication and release.

**Figure 4.**
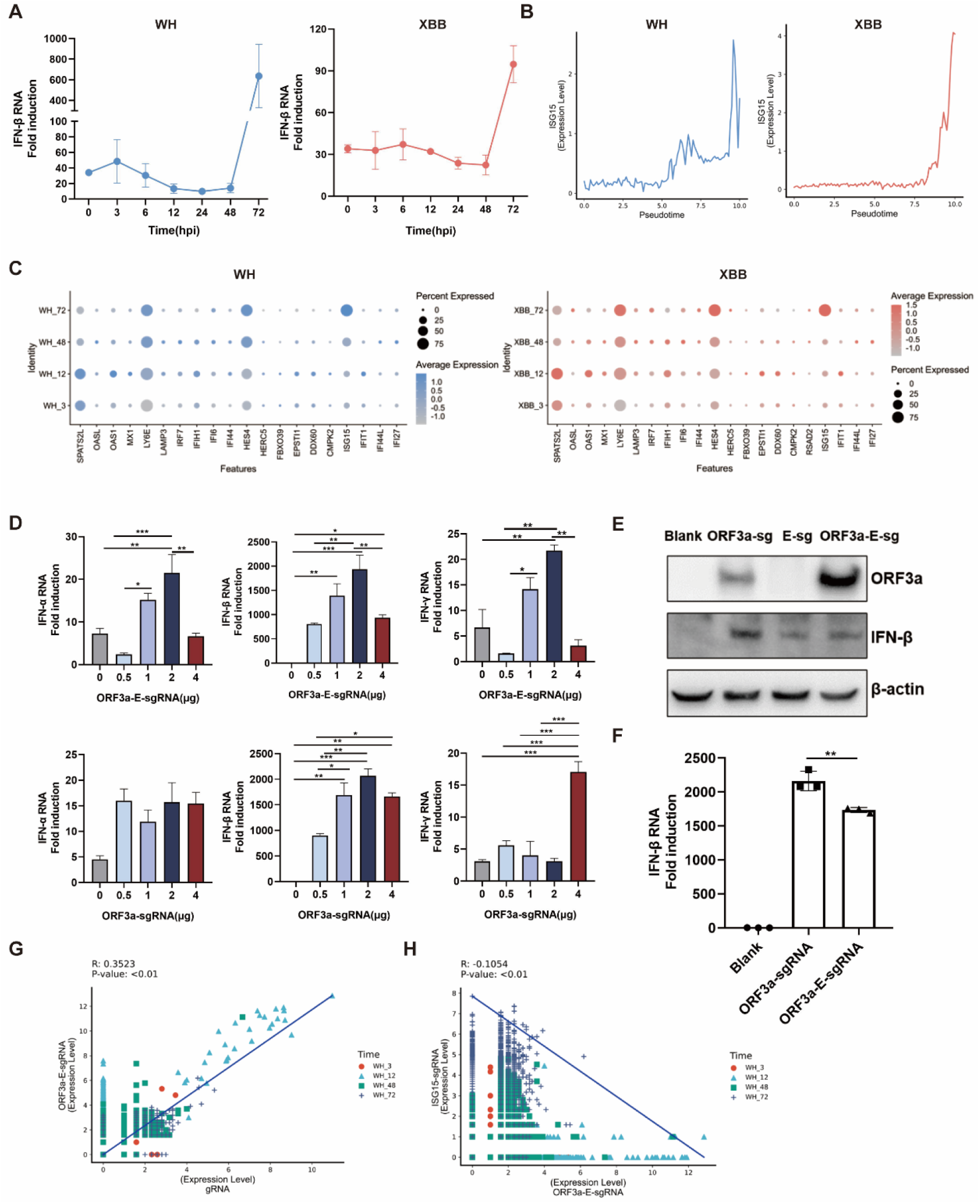
Ability of ORF3a-E-sgRNA to respond during the SARS-CoV-2 life cycle. (A) The expression level of IFN-β in 16HBE cells infected with the Wuhan or XBB strain over the infection period. (B) The expression level of ISG15 in 16HBE cells infected with the Wuhan or XBB strain over pseudotime. (C) Analysis of interferon-stimulated gene sets in the host of 16HBE infected with the Wuhan or XBB strain over the infection period. (D) Measurement of interferon gene levels in 293T cells transfected with different doses of ORF3a-sgRNA or ORF3a-E-sgRNA. (E) Comparison of the protein expression levels of IFN-β in 293T cells transfected with ORF3a-sgRNA or ORF3a-E-sgRNA. (F) Comparison of the transcription levels of IFN-β in 293T cells transfected with ORF3a-sgRNA or ORF3a-E-sgRNA. (G-H) Analysis of the correlation between ORF3a E-sgRNA and genomic RNA(G) and ISG15(H).

## Discussion

The emergence of SARS-CoV-2 variants, driven by factors such as immune pressure from vaccination, antiviral drugs, and environmental changes, has led to varying replication and transmission capacities. A thorough understanding of the molecular mechanisms underlying the viral life cycle is important for characterizing viral properties and developing effective vaccines and antiviral strategies.

The SARS-CoV-2 genome encodes structural and accessory proteins through the transcription of sgRNAs, which are important links between viral replication and host cell function. However, the discontinuous transcription of sgRNAs has resulted in sequence diversity, making them challenging to study. There are few relevant studies on SARS-CoV-2 sgRNA at present. Most current studies on SARS-CoV-2 sgRNAs rely on approaches such as the discontinuous transcription of coronaviruses or the use of sequencing techniques to capture 5’ or 3’ ends. These methods may overlook certain sgRNA types because of the limited understanding of their sequence characteristics.

In this study, we identified a novel SARS-CoV-2 sgRNA containing a fusion of the ORF3a and E genes. This sgRNA, termed ORF3a-E-sgRNA, was detected at multiple time points during SARS-CoV-2 infection of Vero and 16HBE cells.

We believe that this ORF3a-E-sgRNA is produced through a mechanism guided by a transcription-regulating sequence (TRS) in the genome body (TRS-B) and the 5’ leader sequence (TRS-L), but there is a missed template switch between ORF3a and E. Interestingly, the 5’ UTR sequences of ORF3a-sgRNA, E-sgRNA, and ORF3a-E-sgRNA were found to be completely identical.

According to the literature and our sequence analysis of the 5′ UTRs of existing sgRNAs, no consistent variable RNA fragments in the 5′ UTRs of other sgRNAs have been identified. This unique composition of the ORF3a-E-sgRNA may represent a more efficient replication mode adopted by SARS-CoV-2 during evolution, allowing large-scale production and self-optimization of sgRNAs to meet the virus’s replication needs and hijack the host translation machinery. Importantly, our findings suggest that ORF3a-E-sgRNA persists throughout the SARS-CoV-2 life cycle, accounting for a relatively high proportion of sgRNAs. It possesses the ability to strongly recruit ribosomes, evade immune responses, and encode viral proteins, all of which are beneficial for efficient viral replication. Moreover, we detected differential expression of ORF3a-E-sgRNA in cells infected with the Wuhan or XBB SARS-CoV-2 variant, suggesting that ORF3a-E-sgRNA may influence infection outcomes in different viral strains. These changes were accompanied by changes in the endoplasmic reticulum, Golgi apparatus, and cellular energy metabolism.

## Methods

RT−PCR: All primers were synthesized by Qingke Biotechnology (Table S1). Total RNA was extracted from the samples and transcribed via a reverse transcription kit (Takara, 6215A), after which PCR was performed to determine the sequence characteristics of the sgRNA.

In vitro transcription: The target sequence containing the T7 promoter was obtained through PCR amplification (Takara, R045A) and confirmed through sequencing; the sequence was transcribed to mRNA, and a cap and tail (NEB, E2060S) were added, labeled with Cy3(Enzo, ENZ-42505), Cy5 (Enzo, ENZ-42506) and biotin via UTP (Roche, 11388908910).

mRNA transfection: mRNA transfection was carried out via Lipofectamine Messenger MAX mRNA Transfection Reagent (Thermo Fisher, LNRNA001). A total of 125 µl of Opti-MEM was used to dilute 7 µl of transfection reagent and 2 µg of mRNA, after which the solution was incubated for 10 minutes and mixed. The mixture was added to the cells, and the cells were cultured in an incubator.

In situ hybridization: The probe used for hybridization was synthesized by BGI. The probe was denatured before the experiment began, and the cell slides were sequentially placed in gradient ethanol solution for rehydration. Protease K was used to break the cell membrane, gradient ethanol solution was used for dehydration of the slides, the probe was used to label the ORF3a-E-sgRNA at 4°C, and the nuclei were stained with DAPI. Fluorescence images were captured via Leica TCS SP8 laser confocal microscopy.

Flim FRET: Cy3 and Cy5 were incorporated during the mRNA preparation through in vitro transcription and used as donors and acceptors, respectively, for in situ hybridization and fluorescence staining. The fluorescence lifetimes of Cy3 in the presence of Cy3 only and after the addition of Cy5 were measured and recorded via Leica TCS SP8 laser confocal microscopy in FLIM mode.

Immunohistochemistry: Paraffin-embedded tissue slides were dewaxed with xylene, dehydrated with gradient ethanol, washed with distilled water, subjected to antigen retrieval with sodium citrate, incubated overnight with SARS-CoV-2 N antigen and analyzed via a tissue chemistry section scanner.

SARS-CoV-2 infection: Vero, 16HBE and AC16 cells were cultured in 12-well cell culture plates, and when they reached 70%-80% confluence, virus maintenance medium containing 2% FBS was used. A total of 1 × 10^6^ TCID50/ml SARS-CoV-2 virus was used for infection, and cell samples were collected at different times during infection. All experiments involving SARS-CoV-2 were conducted in the biosafety cabinet of the biosafety level III facility of the Institute of Medical Biology, Chinese Academy of Sciences.

Western Blotting: The cell samples obtained from viral infection and cell transfection were added to RIPA protein lysis buffer supplemented with protease inhibitors and then lysed on ice for 30 minutes. The supernatant was harvested after centrifugation at 12000 rpm and 4°C for 15 minutes. The samples were quantified via a BCA protein assay kit and boiled in 1X SDS buffer at 95°C for 10 minutes. The activation of the host ribosomal pathway was analyzed via protein blotting after viral infection and transfection. The samples were separated by SDS−PAGE, and the proteins were transferred to a 0.22 µm PVDF membrane via a membrane transfer instrument (GenScript). The membrane was incubated with the primary antibody (RPS3, abcam, Cat#ab128995; eIF4E, abcam, Cat#ab33768; ORF3a, abcam, Cat#280953; 5-mC, Cat#ab214727; IFN-beta, abcam, Cat#ab275580; beta-actin, genetex, Cat#109639) at 4°C overnight, followed by incubation with the secondary antibody at room temperature for 1 hour. An imaging device was used for band visualization.

Single cell sequencing: RNA was extracted and reverse transcribed from 16HBE cells infected with SARS-CoV-2 at different time points in the BSL3 laboratory, followed by RNA isolation, library construction, and sequencing on Singleron Biotechnologies. Among them, sgRNA is captured specifically by designing probes on the 5’UTR and sgRNA sequences. All RNA seq data were analyzed using R software. the Gene Ontology (GO) was used with the “clusterProfiler” R package 3.16.1. Cell differentiation trajectory was reconstructed with Monocle2. Correlation analysis uses a linear fitting line between ORF3a-E-sgRNA and other sgRNAs or genes to calculate the Pearson correlation coefficient. Cell distribution comparisons between two groups were performed using unpaired two-tailed Wilcoxon rank-sum tests. Comparisons of gene expression or gene signature between two groups of cells were performed using unpaired two-tailed Student’s t test. Comparisons of cell distribution of WH and XBB were performed using paired two-tailed Wilcoxon rank-sum tests. Statistical tests used in figures were shown in figure legends and statistical significance was set at p < 0.05. Exact value of n was shown in the figures and figure legends.

## Quantification and statistical analysis

All statistical analyses were conducted using GraphPad Prism 8 software. Data were presented as mean ± SD. For comparisons between the two groups, parametric Student’s t test was used. When more than two groups were compared, two-way anova test was used. Statistically significant differences are represented in figures as *, **, ***, **** for p values < 0.05, <0.01, <0.001, and <0.0001, respectively.

## Acknowledgments

This work was supported by National Natural Science Foundation of China (32070923), Spring City Plan the High-level Talent Promotion and Training Project of Kunming (2022SCP010) Science and techonology talent and platform plan of Yunnan Province (202305AC160008, 202305AD160006), Yunnan Provincial Biomedical Project (202402AA310018). The funders had no role in the study, design, data collection and analysis, decision to publish, or preparation of the manuscript.

## Author contributions

LD.L and YF.Z designed the study and wrote the manuscript. YF.Z performed the experiments and analyzed the data. HW.Z and J.L provided critical expertise. XL.Z, XZ and JL.L conducted the WB experiment. HW.Z and H.L processed samples for scRNA. X.Z, ZH.Z and KQ.C performed the RT-PCR. YY.L and SS.P performed the IF experiments. HW.Z and HJ.S edited the manuscript, LD.L conceived and supervised the study, interpreted the data, and wrote the manuscript.

